# Plant derived bioactive compounds as potential inhibitors of ZIKA virus: an in silico investigation

**DOI:** 10.1101/2020.11.11.378083

**Authors:** Sheikh Rashel Ahmed, Anik Banik, Sadia Mahjabin Anni, Mohammed Mehadi Hassan Chowdhury

## Abstract

The ZIKA virus has caused a heavy concern everywhere the globe because of its high infectivity and mortality rate. Still, there’s no specific drug or preventive medication to treat ZIKA infection despite comprehensive analysis by the researchers. This study was designed to demonstrate the efficacy of some plant derived bioactive compounds against ZIKV by using both structure and ligand based virtual screening methods. A number of 35 plant metabolites were screened against ZIKA NS2B-NS3 protease (5LC0), Envelop protein (5JHM), Capsid protein (5YGH) and NS5 RNA-dependent RNA polymerase protein (5U04) employing molecular docking approach. Results showed that there have been four metabolites, i.e. Chicoric acid, Luteone, Reserpine and Rosmarinic acid provide highest binding affinity to targeted ZIKV proteins. Crucial binding sites and drug surface hotspots are unraveled for every targeted viral protein. The ADME study showed that neither of the candidate compounds had side effects that would reduce their drug-like properties. As compared, the toxicity pattern analysis has unmasked the non-toxic essence of top drug candidates. The RMSD values of ligand-macromolecule complexes were 2 Å apart from Envelop protein- Chicoric Acid, although the RMSF values showed normal atomic fluctuations within the molecular dynamics analysis, with the exception of Envelop protein- Chicoric Acid. The expected majority of the target class the highest drug candidates is enzyme classes (e.g. protease, hydrolase, phosphatase). In addition, the drug similarity prediction revealed several structural analogs from drugbank such as Isoformononetin (DB04202), Deserpidine (DB01089) and Rescinnamine (DB01180) etc. and these analogs could even be an option for the treatment of ZIKV infections. The study can pave the way for the creation of effective ZIKV medications and preventive measures. We highly recommend further in vivo trials for the experimental validation of our findings.

## Introduction

The Zika virus (ZIKV) belongs to the Flaviviridae family of positive-stranded RNA viruses. In 1947, during experiments on the ecology of yellow fever, ZIKV was first isolated in Uganda from a febrile sentinel monkey by the Rockefeller Foundation (Dick et al., 1952). ZIKV is an arthropod-borne virus that is carried by mosquito vectors and can affect both human and moderately non-human primates extensively (Buechler et al., 2017 and Dick et al., 1952). In Nigeria (Africa), the first ZIKV infection in human is ever reported (Macnarmara, 1954). The first non-African ZIKV strain ‘P6-740’ was isolated in 1966 in Malaysia (Marchette et al., 1969). During an analysis in Brazil, the fatality rate in the case of microcephaly and other severe. The advent of ZIKV was linked with neurological problems such as Guillain-Barré syndrome (GBS) in adults from French Polynesia and microcephaly in neonates from Brazil (Krauer et al., 2017). After ZIKV infection, symptoms appear to develop after an incubation time of 3 to 12 days in symptomatic cases and are reported to be characterized by fever, headaches, rashes, arthralgia, gastrointestinal disease, conjunctivitis, myalgia etc. (Duffy et al., 2009 and Oehler et al., 2014). However, there is no particular therapy, antiviral treatment or vaccine is available for ZIKV infection till now. A variety of possible therapies are being evaluated by researchers (Barrows et al., 2016). However, all of the signs caused for ZIKV infection are treated now, likewise acetaminophen-based treatments for fever and discomfort, an antihistamine for pruritic rash, and fluid consumption (WHO, 2018). Scientific groups, however, are trying to implement effective treatment strategies or to develop effective ZIKA virus drugs. A hope is showing by a FDA approved drug for Hepatitis C named Sofosbuvir that inhibits ZIKV replication targeting virus’s ribonucleic acid, have significantly revealed their anti-viral efficacy in in-vivo trials and a step ahead towards clinical trials (Bullard-Feibelman et al., 2017 and Mumtaz et al., 2017). As a novel drug or vaccine development needs time while alternatively, medicinal plants could be used as early as possible in the manufacture of medicines. The importance of plants due to their medicinal benefit and therapeutic applications as ancient medicines has been documented by many scientific researchers (Azim et al., 2020; Koc et al., 2015; Banik et al., 2020 and Ahmed et al., 2019). Herbal bio-active compounds such as curcumin, a food supplement, have an antiviral activity against the Zika virus (Mounce et al., 2017). Recently, a study found structurally related polyphenols delphinidin (D), and epigallocatechin gallate (EGCG) both have poteintial to impaire infectivity of ZIKV (Vázquez-Calvo *et al*., 2017). Natural plant based bioactive products has the potential to flourish as the basis of modern holistic health care system (Cheuka et al., 2017). The bioactive compound from medicinal plants has the ability to be effective in curing life-threatening diseases such as cancer, Alzheimer’s disease, diabetes, malaria, and heart disease while minimizing the toxicity of medicines (Karimi et al., 2015). Side effects of synthetic products may be bypassed by using of the natural product as a new medication to resist the ZIKV infection and associated complications. Therefore, the purpose of the research was to test some plant-based bio active drug candidate compounds against ZIKV through virtual screening methods and various computational studies (Table 1).

**Table 1:**
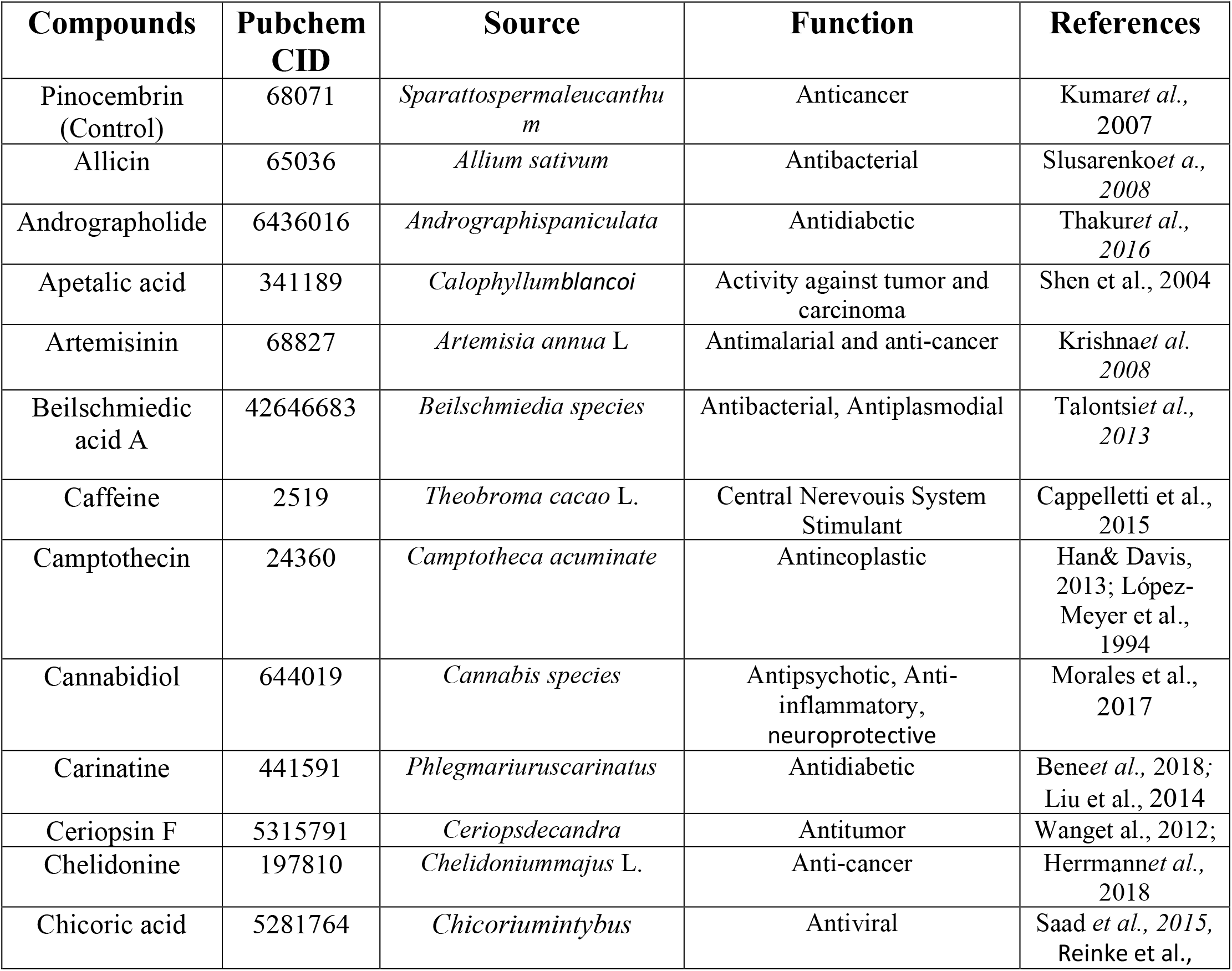
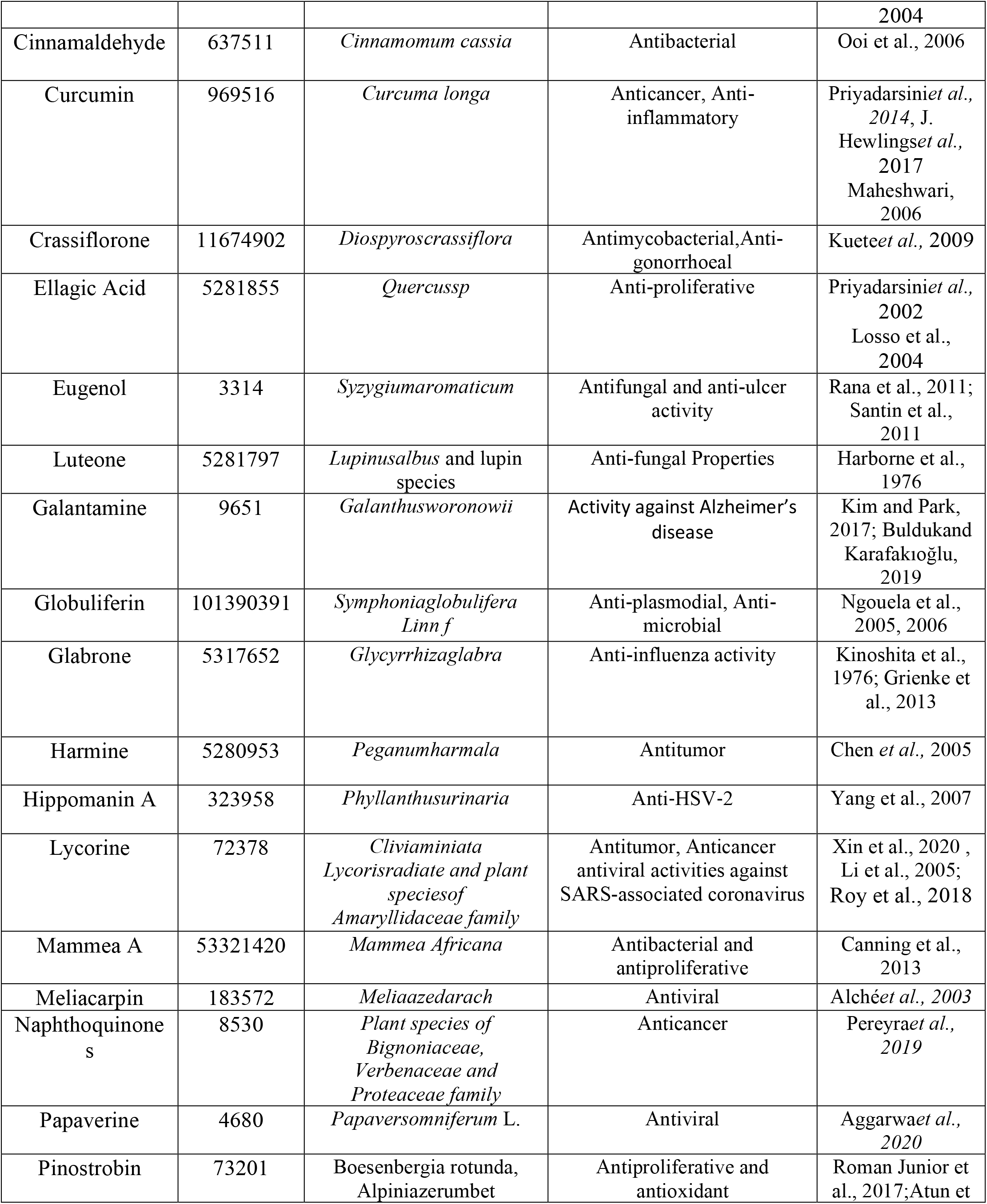
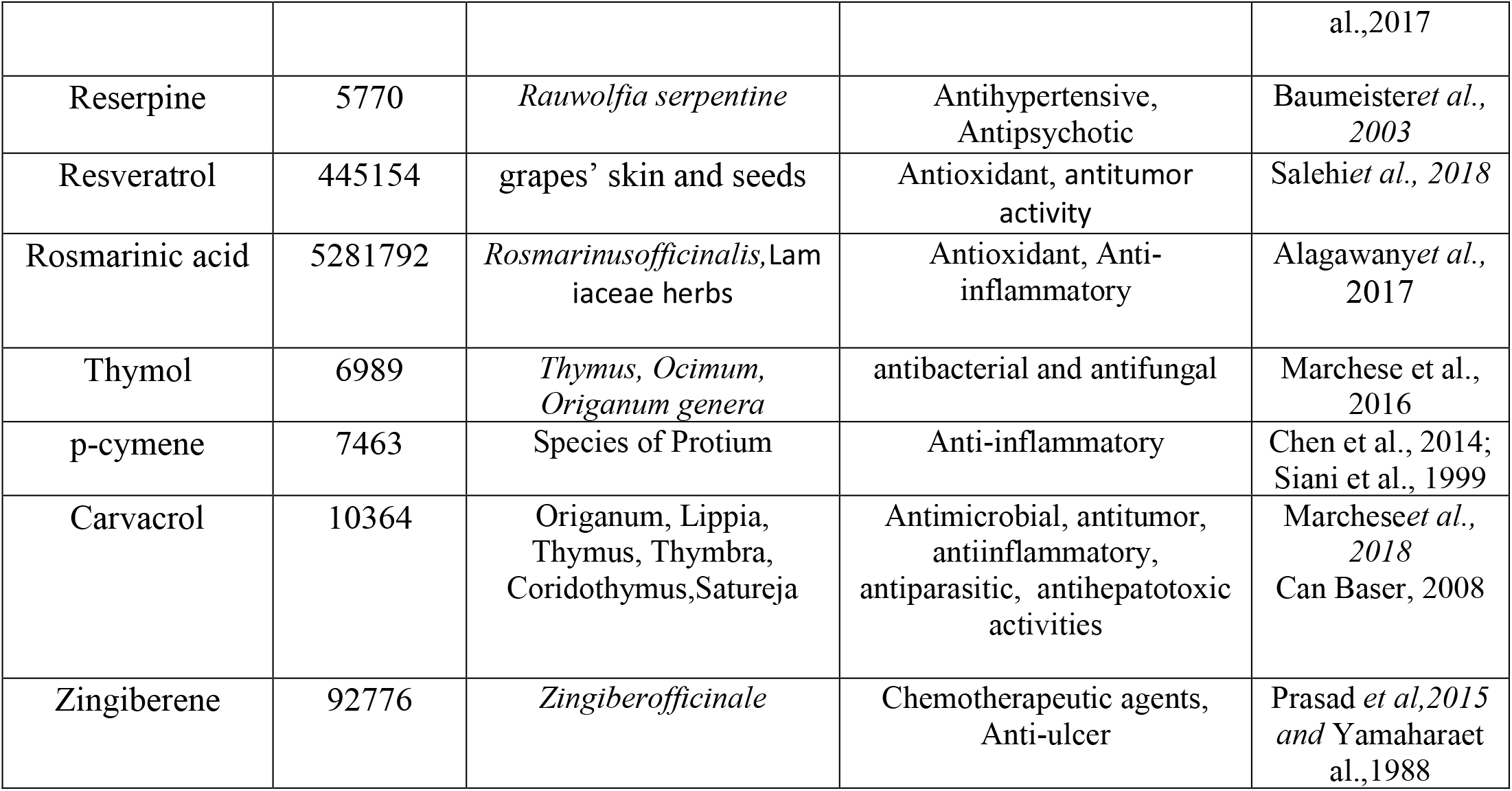
Plant derived compounds’ name, source and their activities.

## 2. Materials and Method

### 2.1. Extraction of Zika virus proteins/protein-domains and plant metabolites

ZIKA virus Envelope protein (5JHM), Capsid Protein (5YGH), NS5 RNA-dependent RNA polymerase (5U04), and NS2B-NS3 protease (5LC0) three dimensional (3D) structure were extracted from the RCSB Protein Data Bank (Rose et al., 2016). Pubchem database was used to retrieve a total 35 metabolites belonging to different class (Table 1) in SDF format (Kim et al., 2016). Further the retrieved SDF structures were converted into the PDB format by Open Babel v2.3 (O’Boyle et al., 2011).

### 2.2. Evaluation of plant metabolites against Zika virus proteins/protein-domains

An effective molecular approach is computer-assisted drug design (Morris et al., 2008). It is ideal for screening therapeutics against particular drug targets of deadly pathogens (Meng et al., 2011). This important approach is used to model the relationship between small ligands and macromolecules, giving a potential chance to a ligand in the way for the discovery of drugs (Kitchen et al., 2004). The binding affinity of 35 plant metabolites with different Zika virus proteins/protein domains (drug targets/macromolecules) were determined by using the PatchDock server (Supplimentary File 1). The PatchDock server complies geometry docking using the shape complementary principals and find out the best patch for the lingand including active sites of the macromolecules (Schneidman-Duhovny et al., 2005 and Banik et al., 2020). The clustering RMSD was set 4.0 and the macromolecule-small ligand type was utilized for docking purpose. Recently, Pinocembrin a compound from honey has been suggested as a Zika virus replication inhibitor by experimental study (Lee et al., 2019) and used as a positive control for this present study. FireDock refinement tool utilized for refinement purpose of the docked complex (Mashiach et al., 2008 and Banik et al., 2020). The ligand bond complexes were visualized by Discovery Studio v3.1 and PyMOL v2.0 (Wang et al., 2015 and DeLano et al., 2002).

### 2.3. Drug surface hotspot analysis and ligand binding pocket prediction

By analyzing the docked complexes with the top metabolites which was Chicoric acid, Luteone, Reserpine and Rosmarinic acid, the drug surface hotspot of Zika virus proteins was evaluated.using several software like Discovery Studio and PyMOL v.2.0 (Wang et al., 2015; DeLano 2002). The comparative structural analysis was employed by utilizing the binding patterns. Moreover, the molecular interaction of pinocembrin (a positive control) with the studied proteins was also analyzed.

### 2.4. Molecular Dynamics Study

By performing normal mode analysis (NMA) via iMODS server (http://imods.chaconlab.org), the conformational stability of the ligand-protein complex interactions was evaluated using various properties like deformability and eigen-values of the protein-ligand interactions (López-Blanco et al., 2014). Further the complexes, the root mean square deviation (RMSD), B factor analysis and the root-mean-square fluctuation (RMSF) were measured and analyzed using LARMD server ((http://chemyang.ccnu.edu.cn/ccb/server/LARMD), uses AMBER16 program from the trajectory with a time interval per picosecond (Roe & Cheatham, 2013; Yang et al., 2019; Banik et al; 2020).

### 2.5. Analysis of top metabolite’s Drug profile

The drug levels and kinetics of drug delivery to the tissues within an organism are determined by four key factors which are absorption, distribution, metabolism and excretion (ADME). The pharmacological activity efficacy was influenced by these parameters (Balani et al., 2005). The SwissADME server was utilized to explore the ADME properties of top four screened metabolites (Daina et al., 2017). The toxicity pattern of these top four metabolites was assessed by pkCSM server (Pires et al., 2015).This server predicts several toxic parameters like LOAEL, LD_50_ etc.

### 2.6. Prediction of drug targets and available drug molecules from DrugBank

To estimate the future macromolecular targets of expected drug candidates, an online based sever SwissTargetPrediction was used (Daina et al., 2019; Rahman et al., 2020). Based on a combination of 2D and 3D similarity, the server estimates a library of 3,70000 recognized bioactive compounds on around 3000 proteins. Moreover, on the basis on homology screening of expected top drug candidates, SwissSimilarity web tools were used to identify possible existing drug molecules that can be repurposed against Zika Virus. The server utilized ligand-based virtual screening of several small molecule libraries to use various approaches including electroshape, spectrophores, FP2 fingerprints, and align-ITI to find approved, experimental, or commercially available drugs from DrugBank (Zoete et al., 2016).

## 3. Results

### 3.1. Evaluation of plant metabolites against ZIKV

The retrieved structure of ZIKA virus crucial protein/protein domain (macromolecule) and plant metabolites (ligands) were optimized and placed for molecular docking stimulation to predict the binding affinity between the macromolecules and ligands which were previously mentioned above (Supplementary File 1). The top ranking of metabolite was done on the basis of global binding energy and the fin dings reveals that the top four scorers (metabolites) were the same in terms of minimum binding energy for each of the macromolecules (Table 2 and Figure 2). In each case, Chicoric acid, Luteone, Reserpine, and Rosmarinic acid display the best binding interactions with four studied ZIKA virus macromolecules (Figure 3). Moreover, Luteone exhibited the highest binding affinity with Capsid Protein (−52.92 kcal/mol), Chicoric acid with Envelope protein (−58.26 kcal/mol), Reserpine with ZIKA virus NS2B-NS3 protease (61.45 kcal/mol) and finally, Rosmarinic acid shows best binding affinity with NS5 RNA-dependent RNA polymerase protein (51.26 kcal/mol) (Figure 2). In additionally, these top predicted 4 metabolites shows very similar binding affinity against among targeted four macromolecules (Figure 2 and Table 2).

**Table 2:**
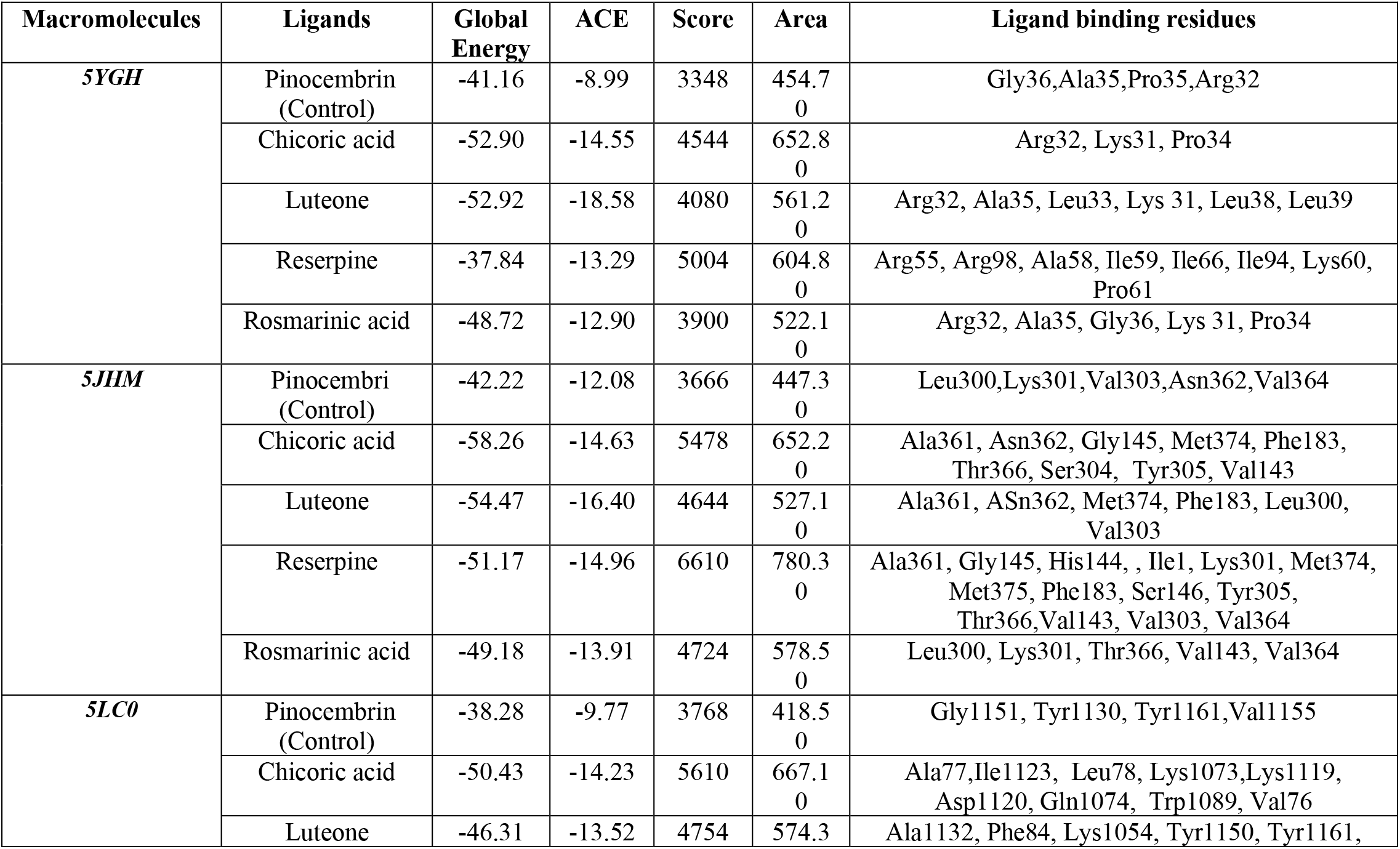
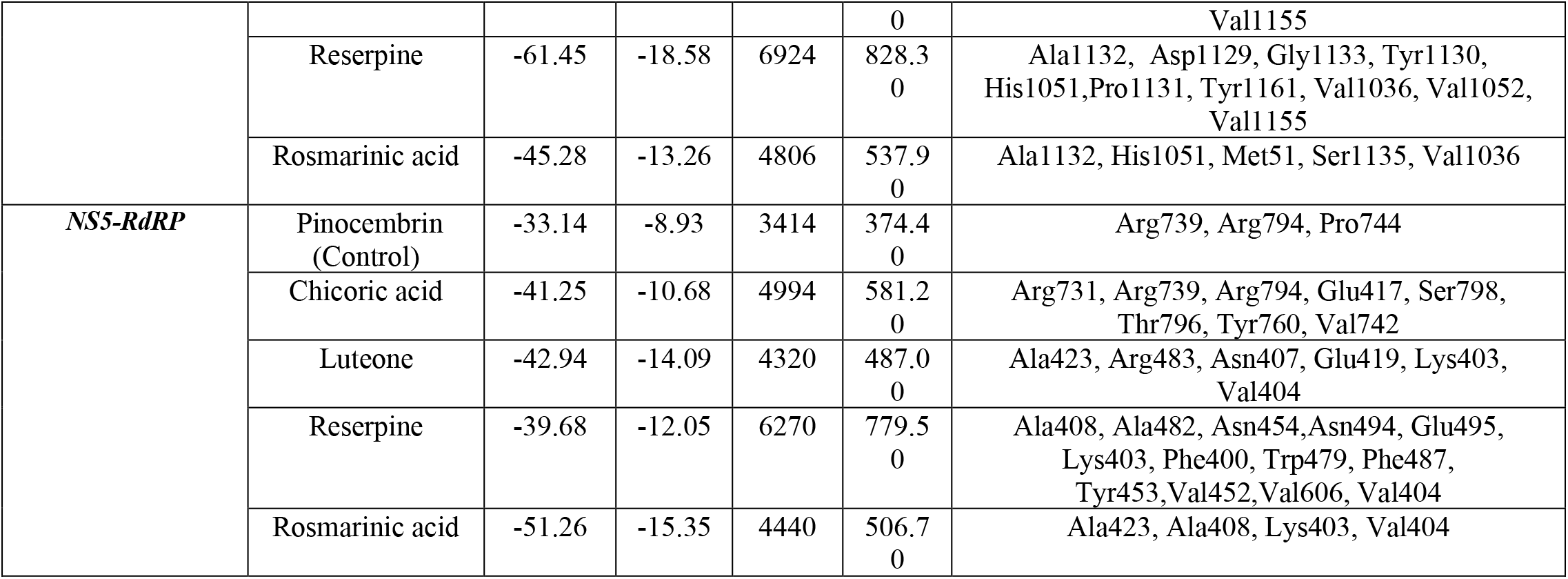
Analysis of global binding energy and interaction sites of the top screened candidates.

**Figure 1:**
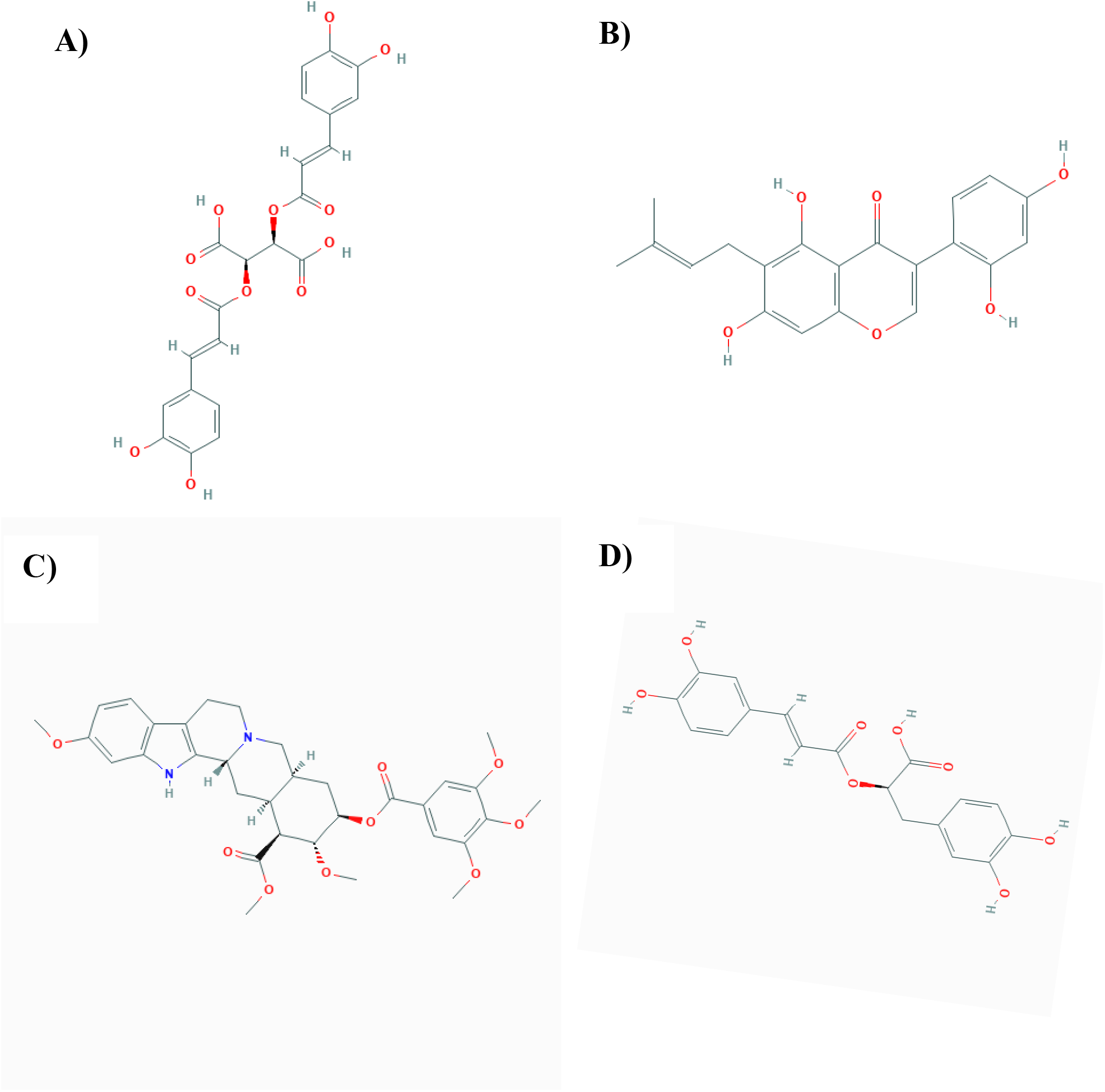
Chemical Structure of Chicoric acid A), Luteone B), Reserpine C), Rosmarinic acid D).

**Figure 2:**
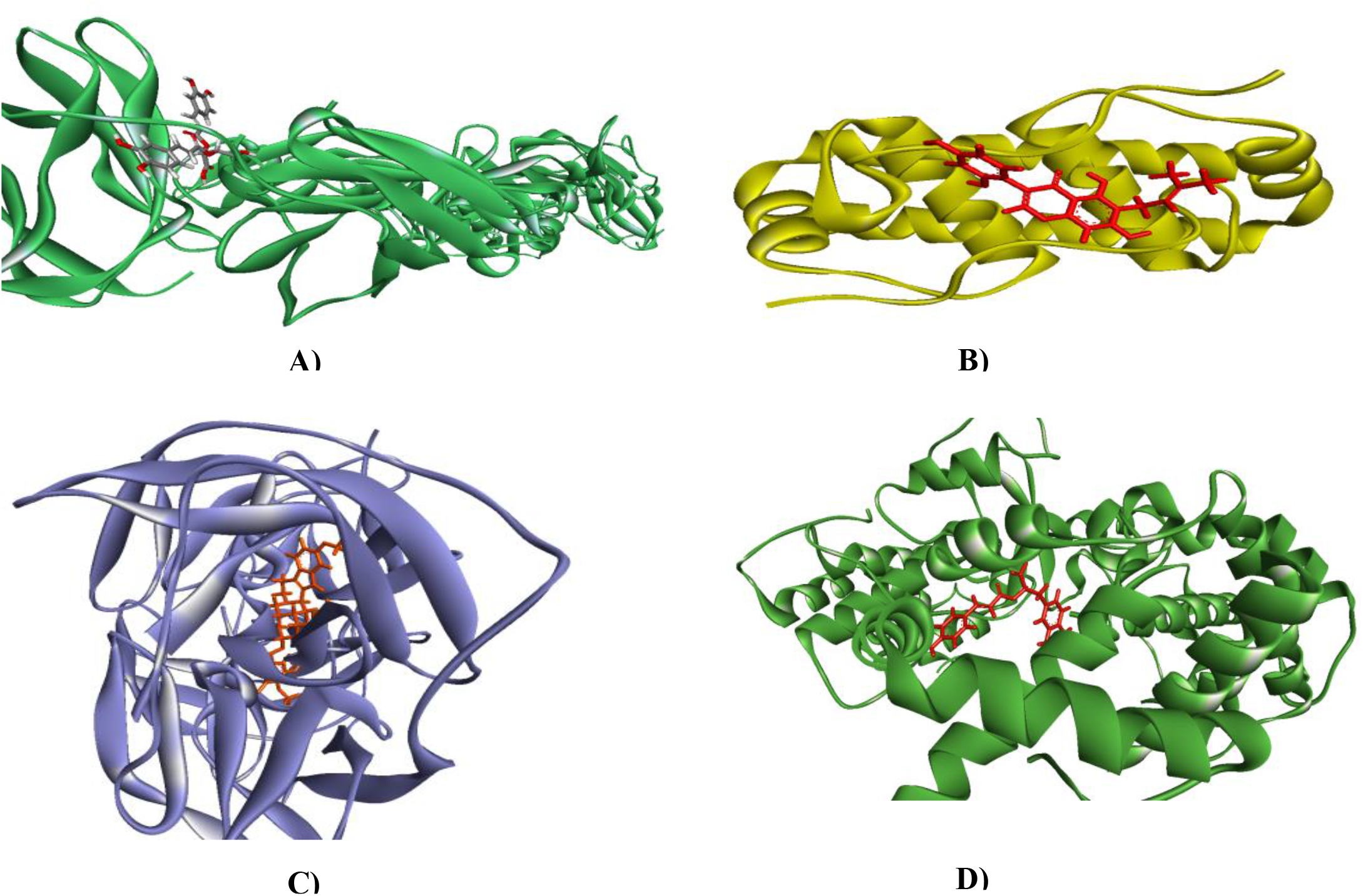
Molecular interactions of macromolecules against ligands; Chicoric Acid A), Luteone B), Reserpine C) and Rosmarinic acid D).

**Figure 3:**
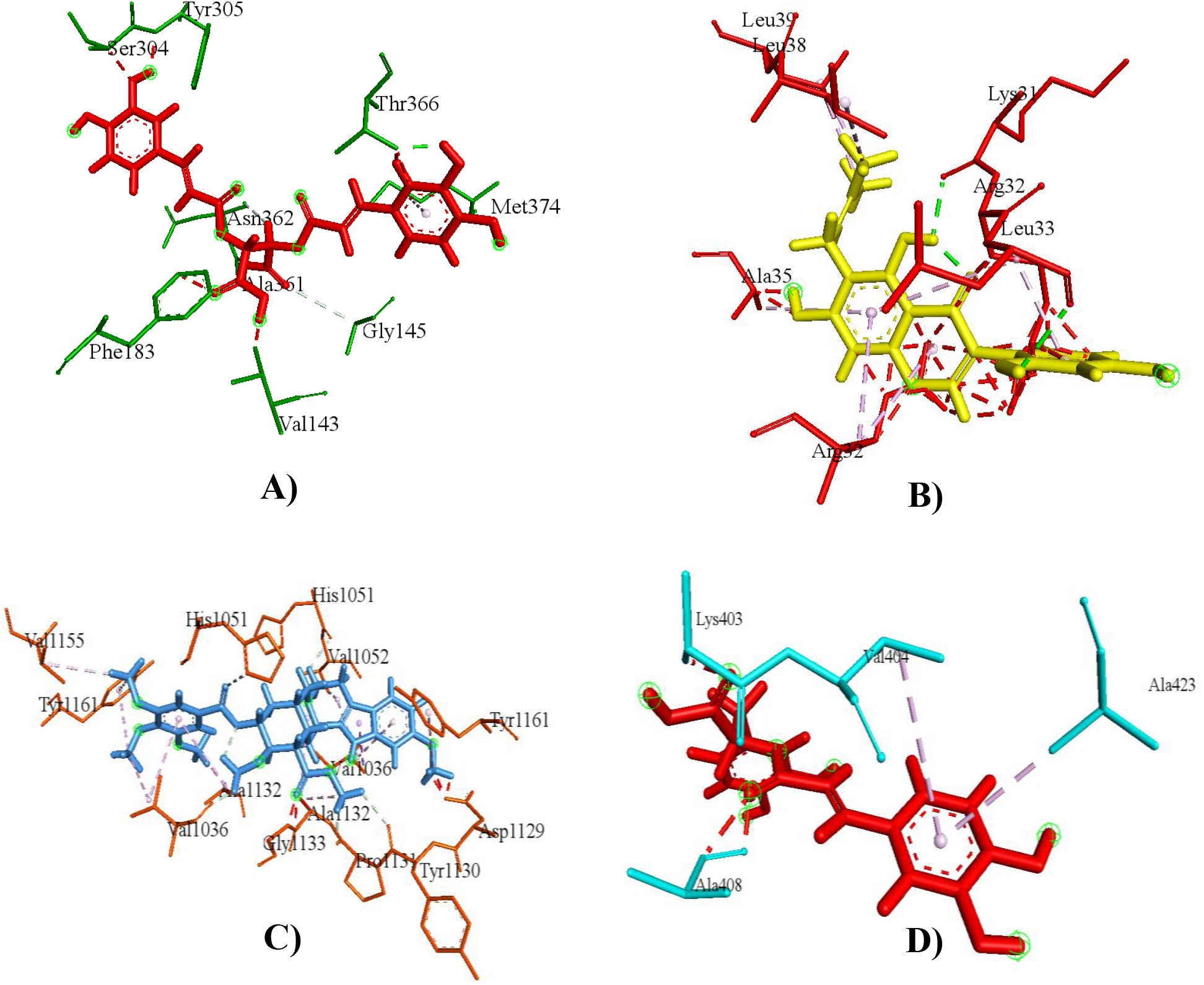
Ligand-Amino acid interaction mode of Chicoric acid with ZIKV Envelope protein A),Luteone with ZIKV Capsid Protein, Reserpine with ZIKV NS2B-NS3 protease C) and ZIKV NS5 RNA-dependent RNA polymerase protein with rosmarinic acid D).

### 3.2. Analysis of drug surface hotspot and ligand binding pocket prediction

To unravel the drug surface hotspot of the studied ZIKA proteins, the structural conformation of the docked complex was analyzed. The ligand binding pattern and residues interacting with their respective positions have been investigated (Table 2). Amino acids from 31 to 61 positions were crucial for the binding interactions of ZIKA Capsid protein (5YGH). Where capsid protein Arg32, Ala35, Leu33, Lys31 amino acid was involved in most cases to formed the docked complex with phyto-ligands. The ligands shows better binding affinity for 143-145, 301-303 and 361-374 regions for envelop protein (5JHM) and most dominant amino acids for interactions with the ligands were Leu300,Lys301, Val303, Val364,Ala361, Asn362, Thr366, Val143, Gly 145.The crucial binding sites for ZIKA virus NS2B-NS3 protease (5LC0) remains into the amino acid 1132-1162 region. The Tyr1161, Val1155, Tyr1130, Ala1132 acid binding sites are found maximum times for the docked complexes. The protein amino acid 403-423 regions were identified as top surface hotspots for NS5 RNA-dependent RNA polymerase protein (5U04); among binding residues Ala423, Ala408, Lys403 were found most of the cases.

### 3.3. Molecular Dynamics stimulation

The deformation of the structures was primarily based on the hinges of the structure. The hinges found in the entire structure are not critical and are stable (Figure 4A, 5A, 6A, 7A). The study of the B factor revealed that there were no significant fluctuations, indicating very less loop numbers (Figure 5B, 6B, 7B) except Envelope-Chicoric acid complex (Figure 4B). The Eigen values for the complexes were higher and the structure was compact and it demonstrated its resistance to deformation. The Eigen values 7.60335 × 10^−5^ for Capsid-Luteone complex, 2.769616 × 10^−5^ (Figure 5C) for Envelope-Chicoric acid complex (Figure4C), and comparatively higher 5.563876 × 10-4 for NS2B-NS3 Protease-Reserpine (Figure 6C) and 7.483752 × 10^−5^ for NS5 RNA-dependent RNA polymerase protein-Rosmarinic acid complex, respectively (Figure7C). The complex RMSD value for all studied complex was less than 2 Å in studied complexes (Figure 5B, 6B, 7B) except Envelope-Chicoric acid complex (Figure 4D). The NS5 RNA-dependent RNA polymerase protein-Rosamarinic complex and Capsid-Luteone shows equilibrium state (Figure 5D, 7D). All the structure didn’t show any major repulsion. For all the complexes the RMSF values showed a regular pattern of atomic fluctuations during the molecular dynamics stimulation (Figure 5E, 6E, 7E) apart from the Envelope-Chicoric acid complex it shows the higher fluctuations may be because of loop region (Figure 4E).

**Figure 4:**
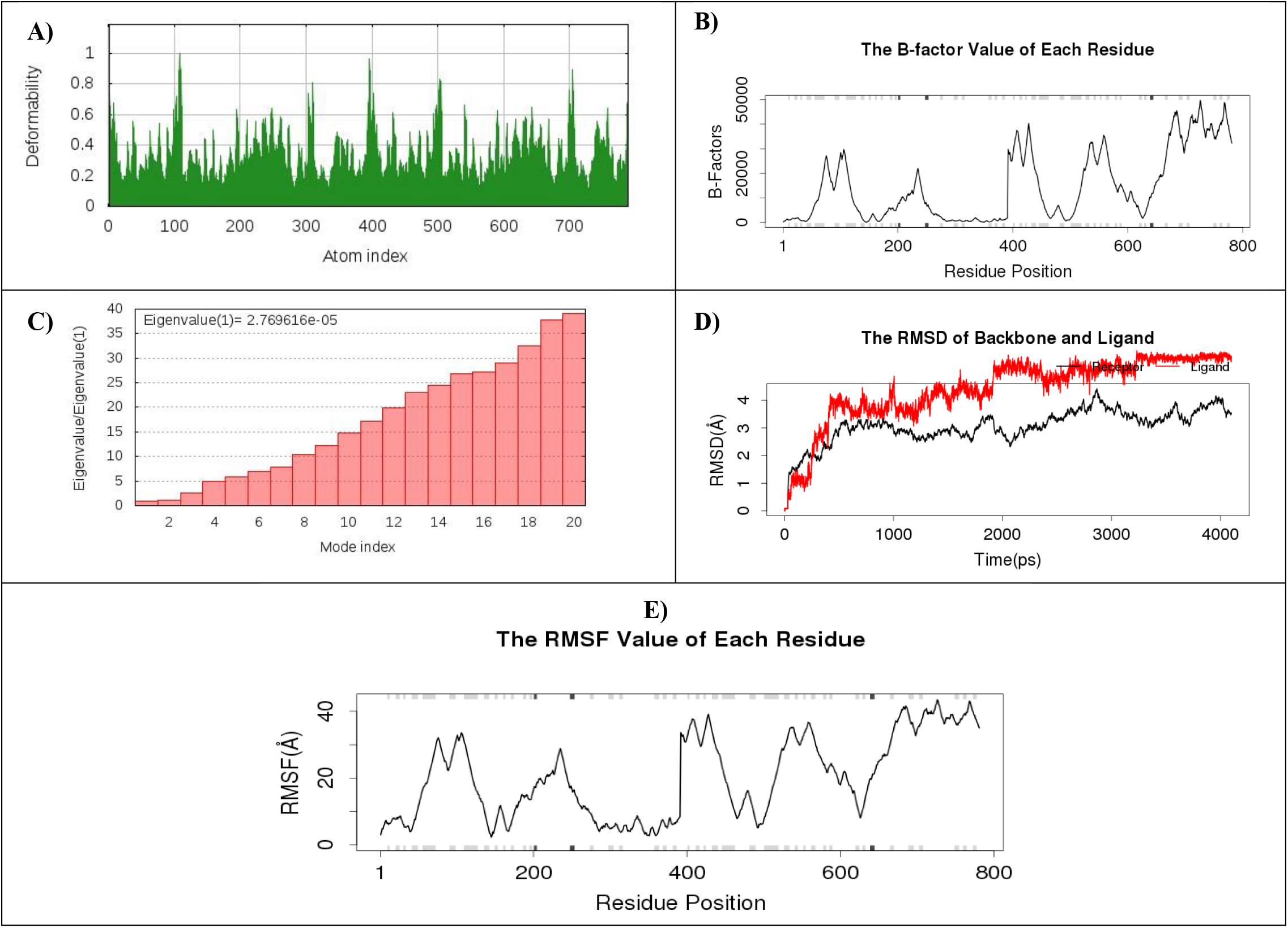
Molecular Dynamics of Envelope Protein-Chichoric acid complex: Deformability analysis A); B factor Analysis B); Eigen value C); RMSD plot D); RMSF value E).

**Figure 5:**
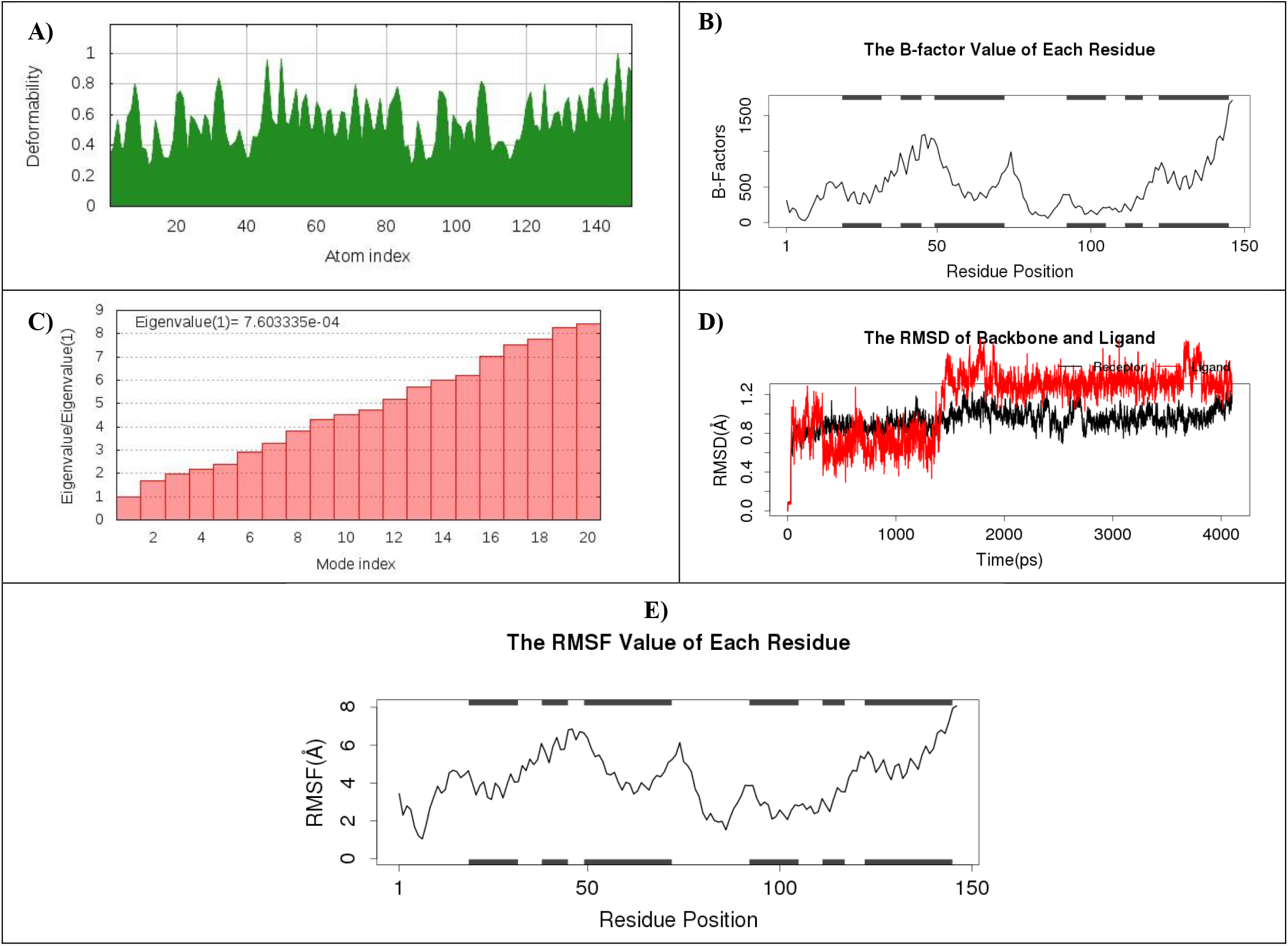
Molecular Dynamics of Capsid Protein-Luteone complex: Deformability analysis A); B factor Analysis B); Eigen value C); RMSD plot D); RMSF value E).

**Figure 6:**
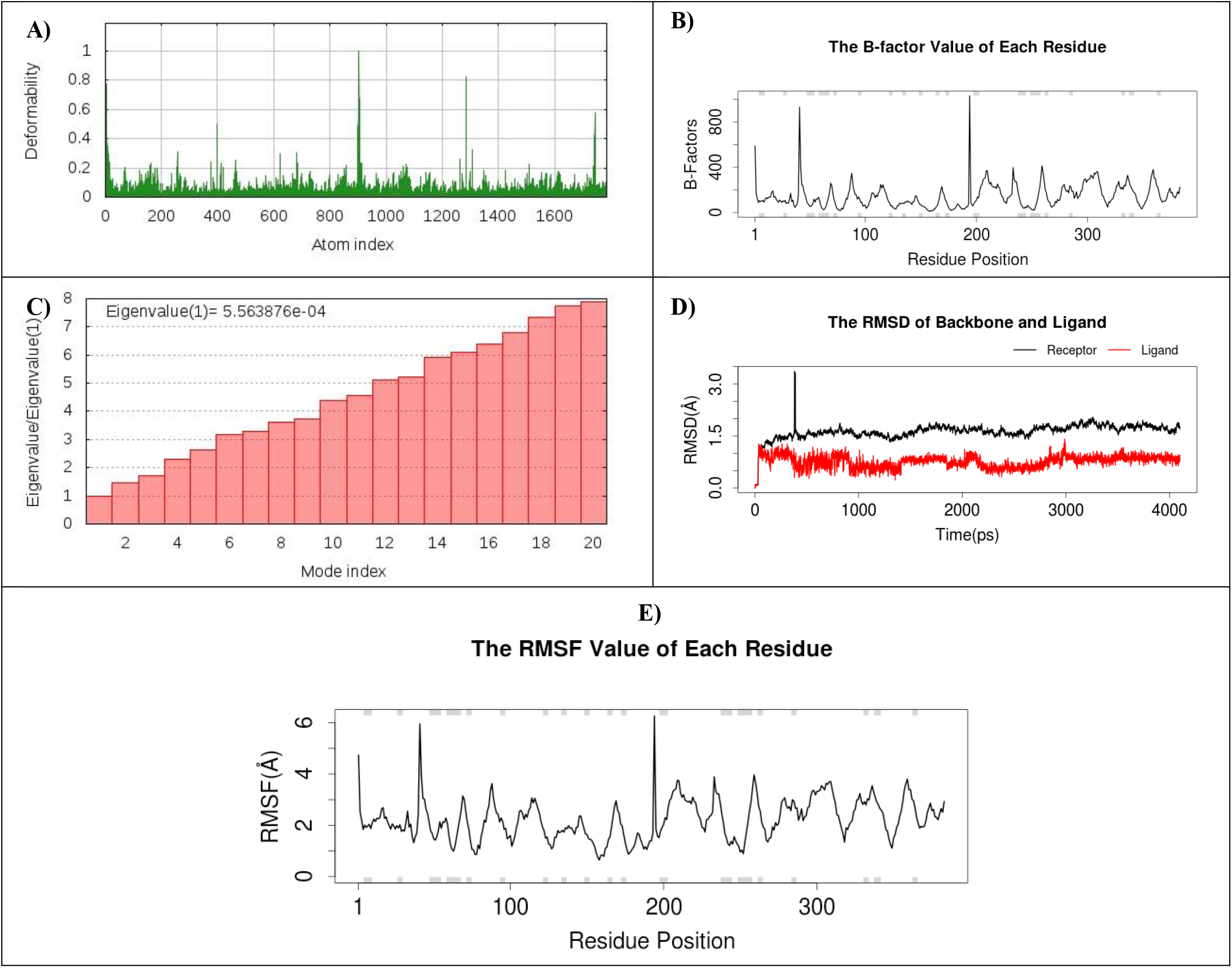
Molecular Dynamics of NS2B-NS3 Protease-Reserpine complex: Deformability analysis A); B factor Analysis B); Eigen value C); RMSD plot D); RMSF value E).

**Figure 7:**
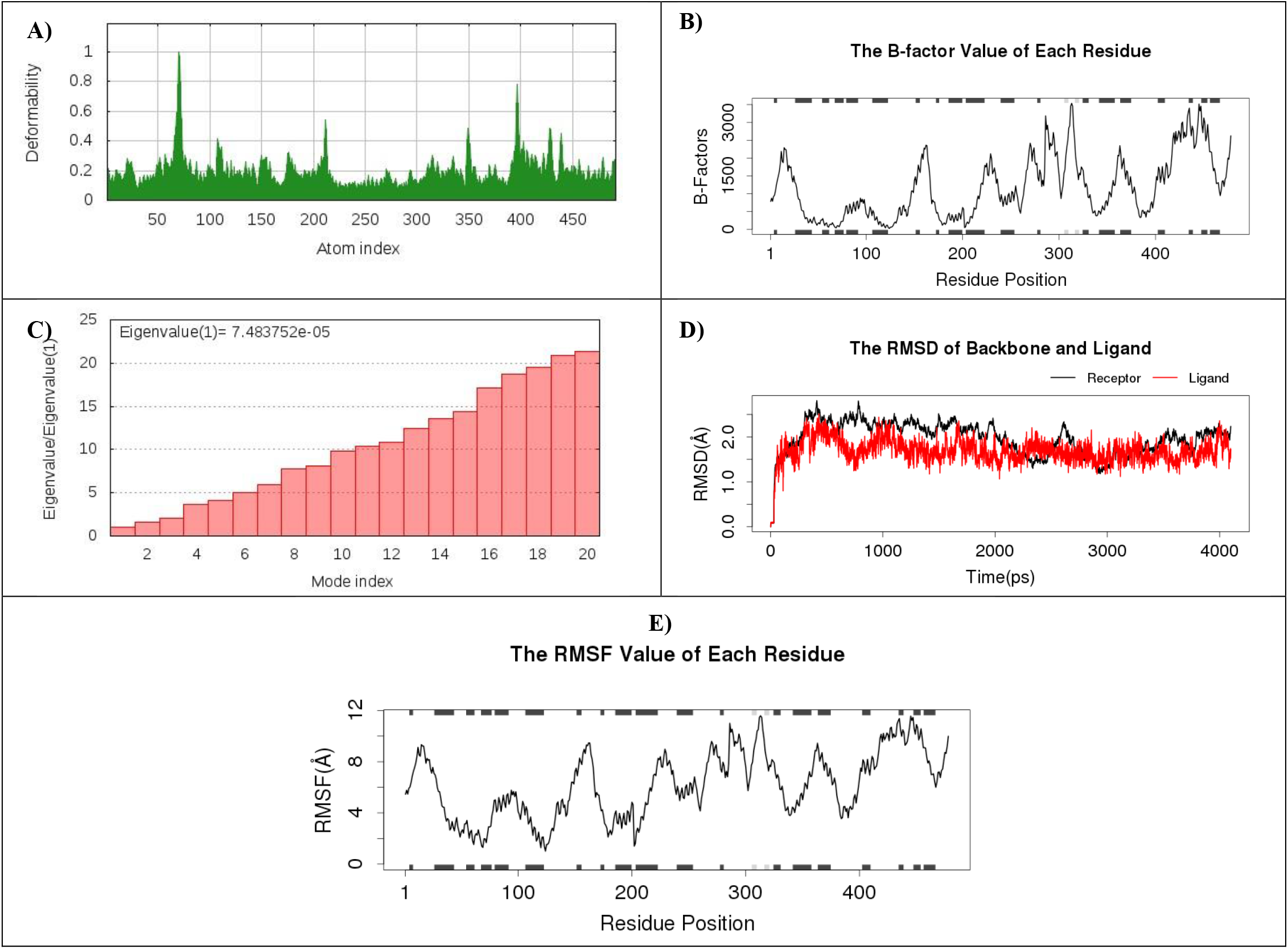
Molecular Dynamics of NS5 RNA-dependent RNA polymerase protein-Rosamarinic complex: Deformability analysis A); B factor Analysis B); Eigen value C); RMSD plot D); RMSF value E).

### 3.4. ADME analysis of top drug candidates

In order to test their drug profiles, various ADME properties, i.e. pharmacokinetics, lipophilicity, water solubility, physicochemical parameters, medicinal chemistry were calculated for the top drug candidates (Figure 8 and Table 3). Examination of inhibition results with various isoforms of CYP (CYP2D6, CYP1A2, CYP2C19, CYP2C9, CYP3A4) reveals that among four, only luteone has the potential to associate with any isoforms of P450 cytochromes. GI absorption was found to be higher for Luteone and Reserpine, while chicric acid and rosmarinic acid were found to be lower. In addition, we used the BOILED-Egg model to measure the blood-brain barrier (BBB) permeation, which showed no BBB permeating among the top drug candidates tested. Each candidate was found water soluble (Table 3).

**Table 3:**
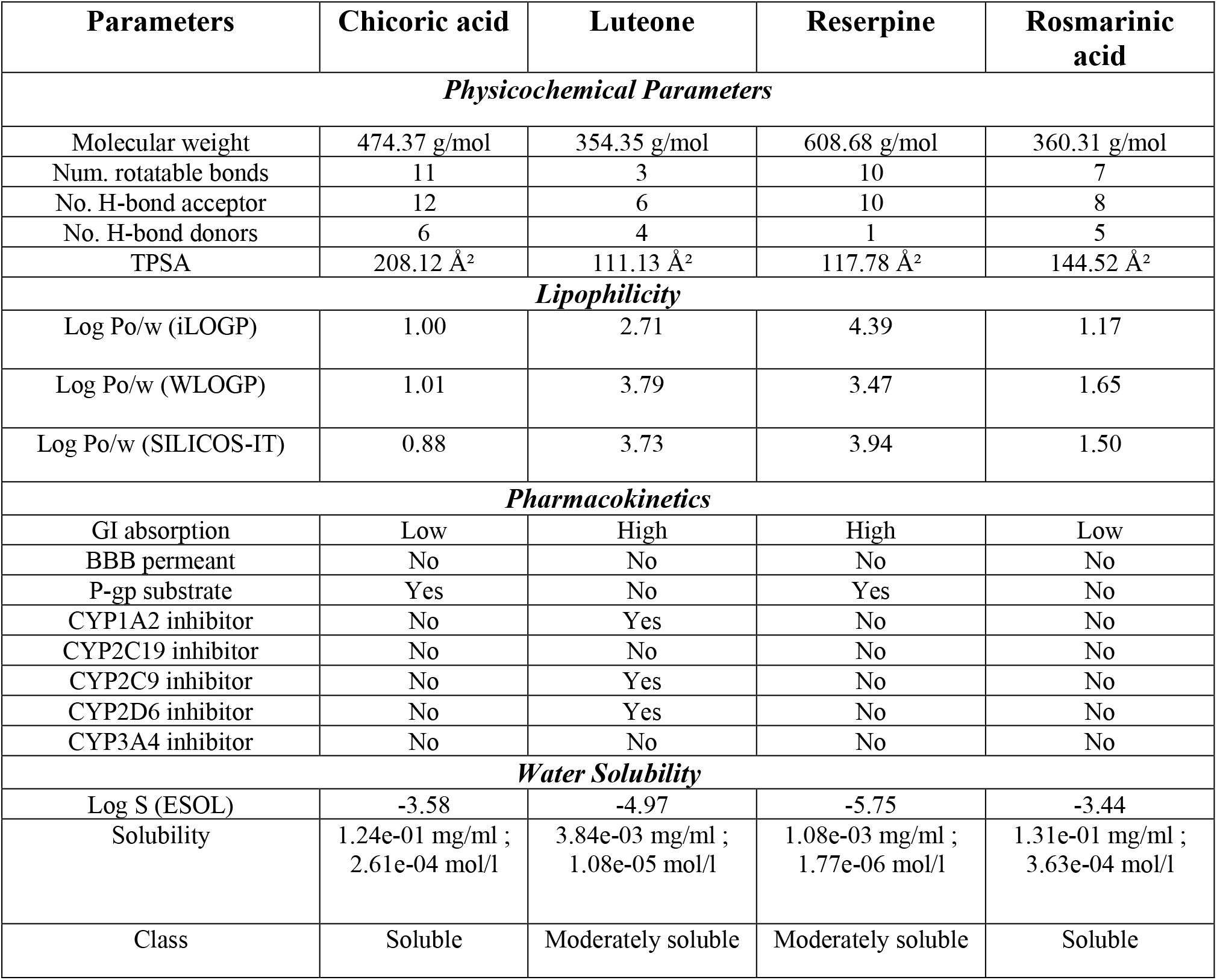
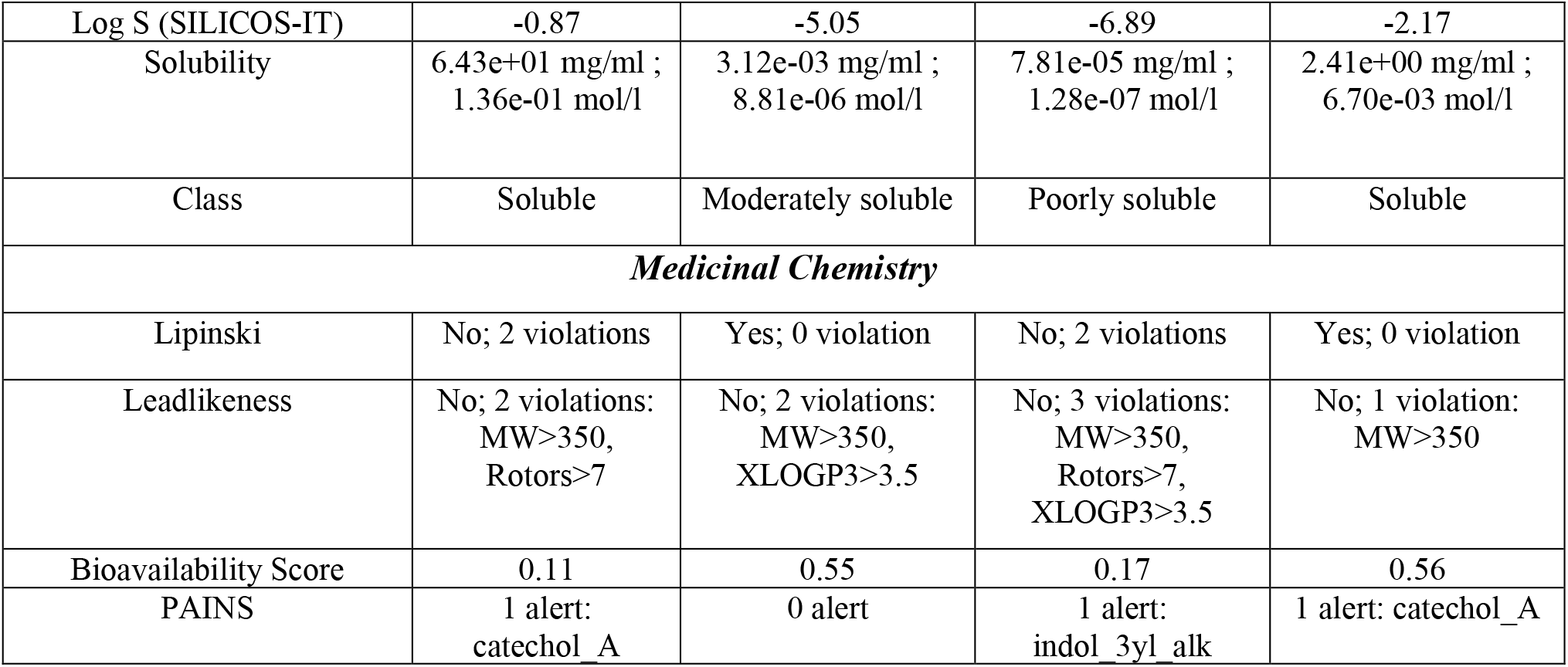
Drug profile and ADME analysis of the top four metabolites.

**Figure 8:**
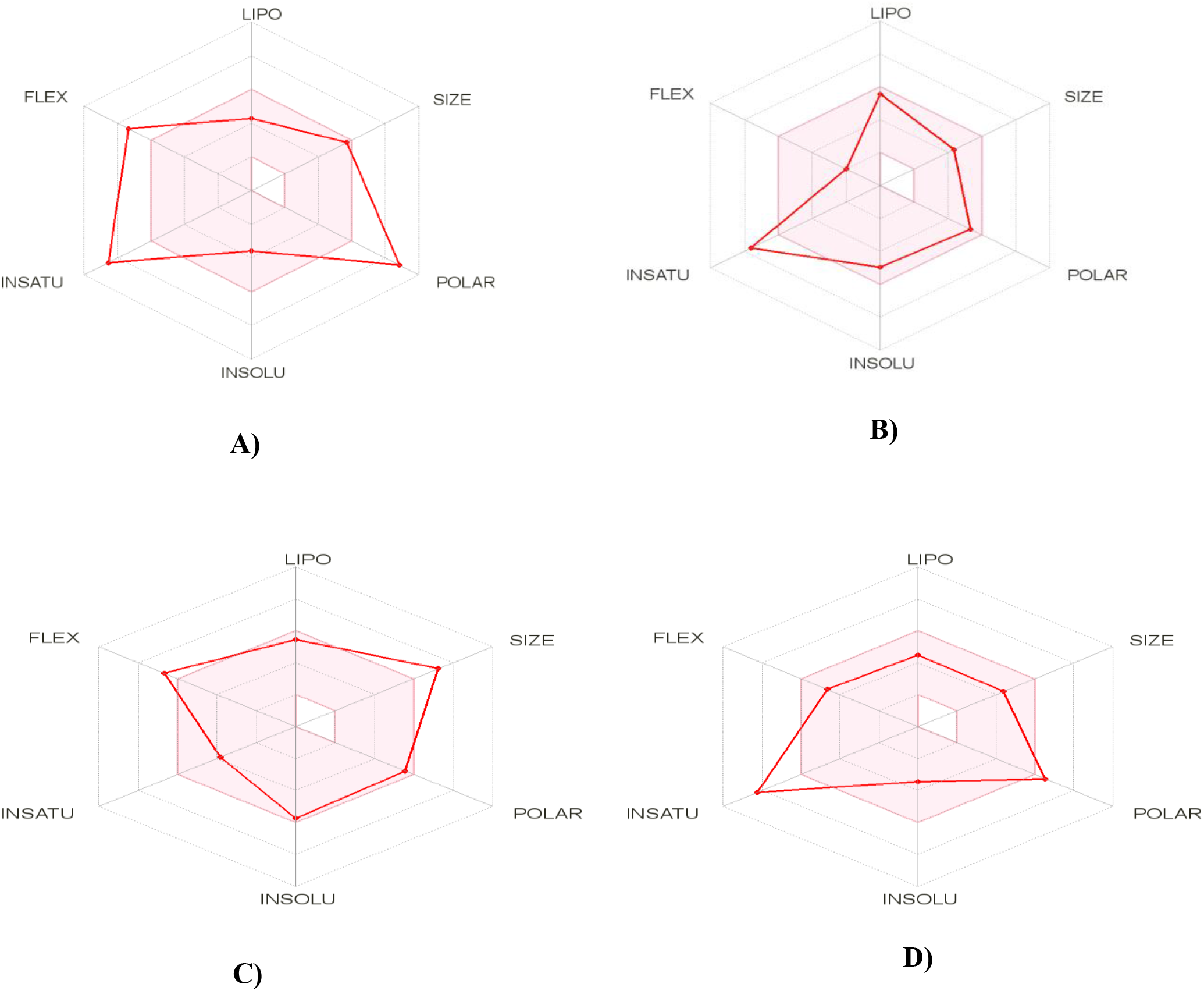
ADME analysis of top four metabolites; Chicoric acid A), Luteone B), Reserpine C), and Rosmarinic acid D).

### 3.5. Toxicity pattern analysis of top drug candidates

The toxicity pattern of top screened candidates states four of them were negative to, carcinogenicity, skin sensitization, Eye corrosion, AMES toxicity. While only luteone may responsible for eye irritation. The all candidates were not responsible for hepatotoxicity and *T. pyriformis* toxicity. The LD_50_ for chicoric acid, luteone, reserpine and rosmarinic acid was 2.445, 2.504, 2.958 and 2.811, respectively (Table 4).

**Table 4:** Toxicity properties analysis of top screened drug candidates.

| Toxicity properties | Chicoric acid | Luteone | Reserpine | Rosmarinic acid |
| --- | --- | --- | --- | --- |
| Carcinogenicity | No | No | No | No |
| Eye corrosion | No | No | No | No |
| Eye irritation | No | Yes | No | No |
| AMES toxicity | No | No | No | No |
| Max. tolerated dose (human) | 0.225 | 0.581 | -0.133 | 0.152 |
| hERG I inhibitor | No | No | No | No |
| Oral Rat Acute Toxicity (LD50) | 2.445 | 2.504 | 2.958 | 2.811 |
| Oral Rat Chronic Toxicity (LOAEL) | 4.005 | 2.191 | 1.158 | 2.907 |
| Hepatotoxicity | No | No | No | No |
| Skin Sensitisation | No | No | No | No |
| *T.Pyriformis* toxicity | 0.285 | 0.319 | 0.286 | 0.302 |

### 3.6. Prediction of drug targets and available drug molecules from DrugBank

The predicted majority of the target class belonged Aldose reductase, Matrix metalloproteinase, Muopioid receptor, Transthyretin etc. (Figure 9 and Table 5). The ligand structural based virtual screening was performed to predict similar bioactive structural compound from DrugBank. The result shows Reserpine similar to Deserpidine (DB01089), Rescinnamine (DB01180) weith higher score i.e. 0.978, 0.925, respectively. The rosmarinic acid and chicoric acid both has structural similarity with 2-deoxy-3,4-bis-o-[3-(4-hydroxyphenyl)propanoyl]-l-threo-pentaric acid (DB08322). Finally, luteone shows a structural similaraity with antiapoptosis drug i.e. Isoformononetin (DB04202) (Table 6). These similar drugs may also used to fight with ZIKA virus needs further *in vivo* trials.

**Table 5:**
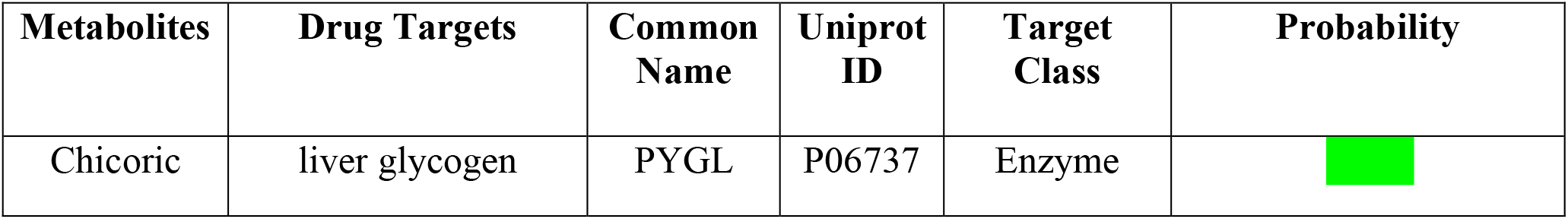
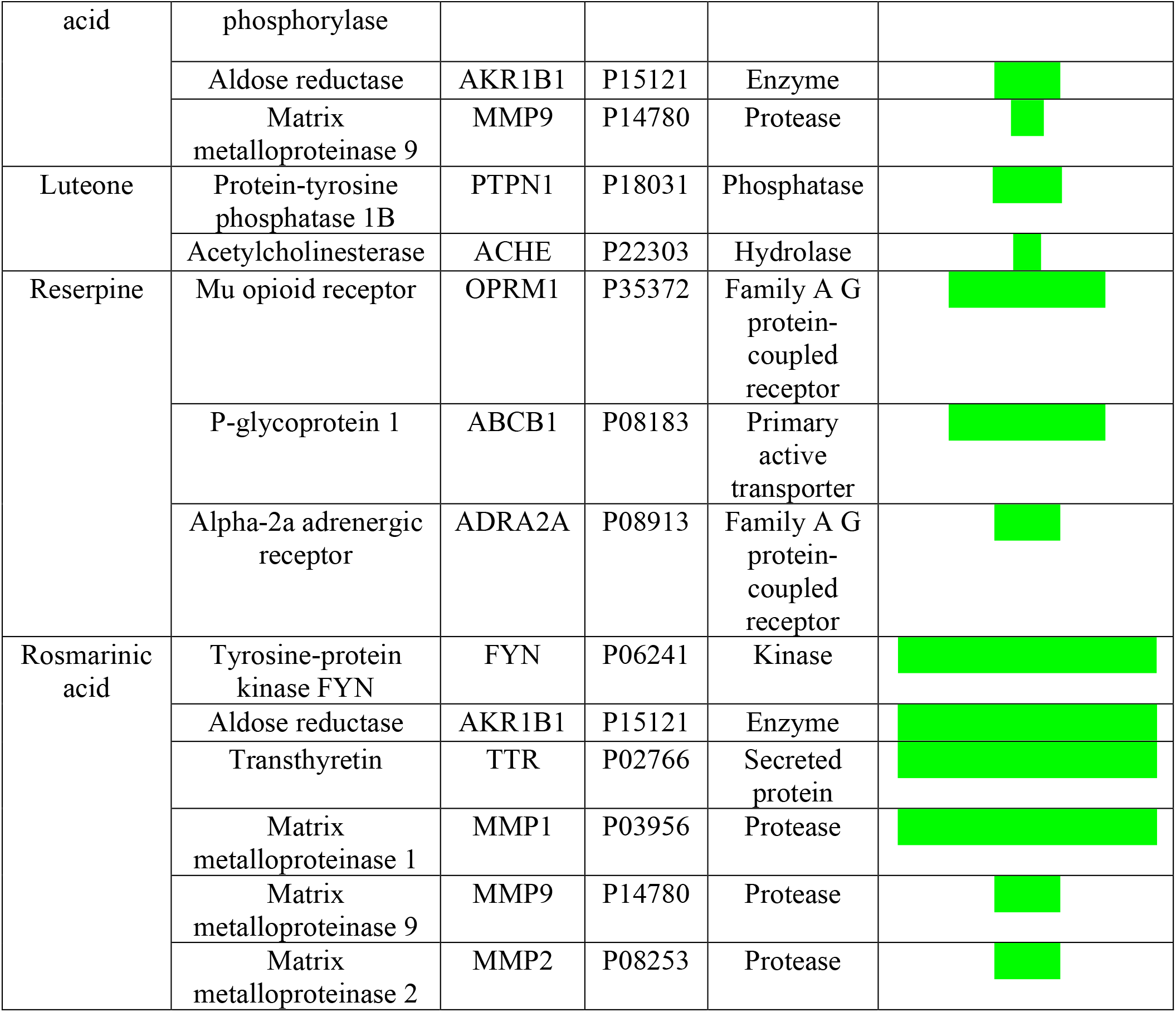
Predicted drug targets of top screened candidates.

**Table 6:**
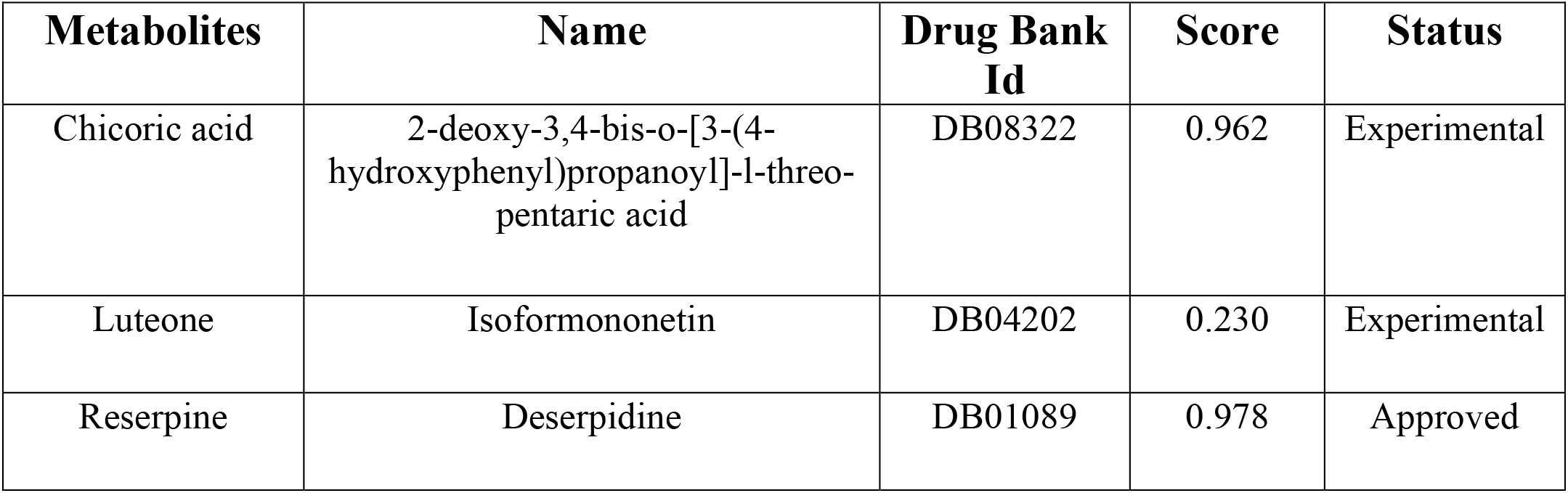
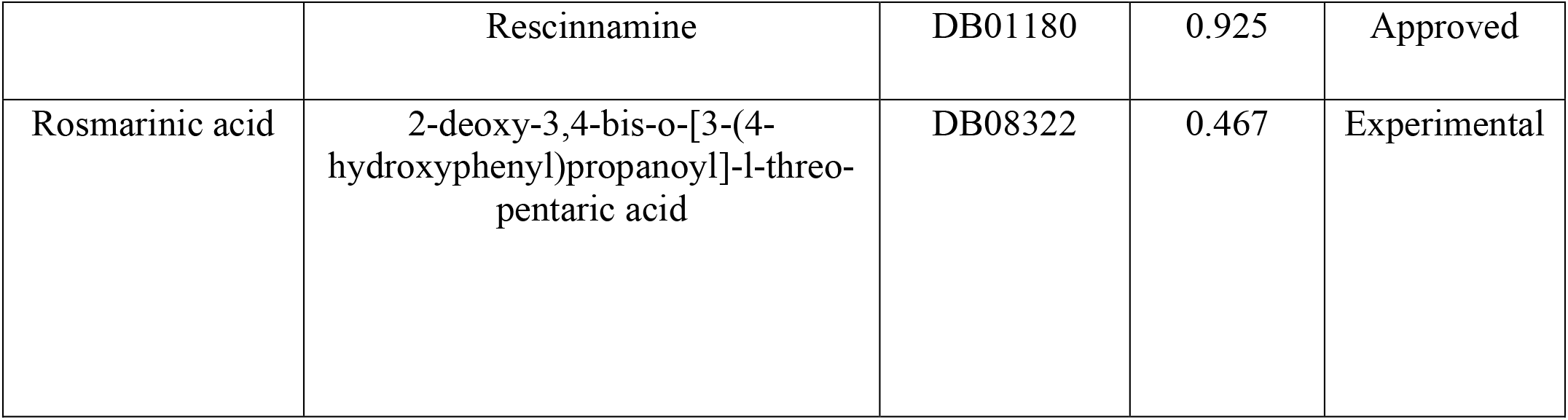
Predicted Structural analog molecules from Drugbank.

**Figure 9:**
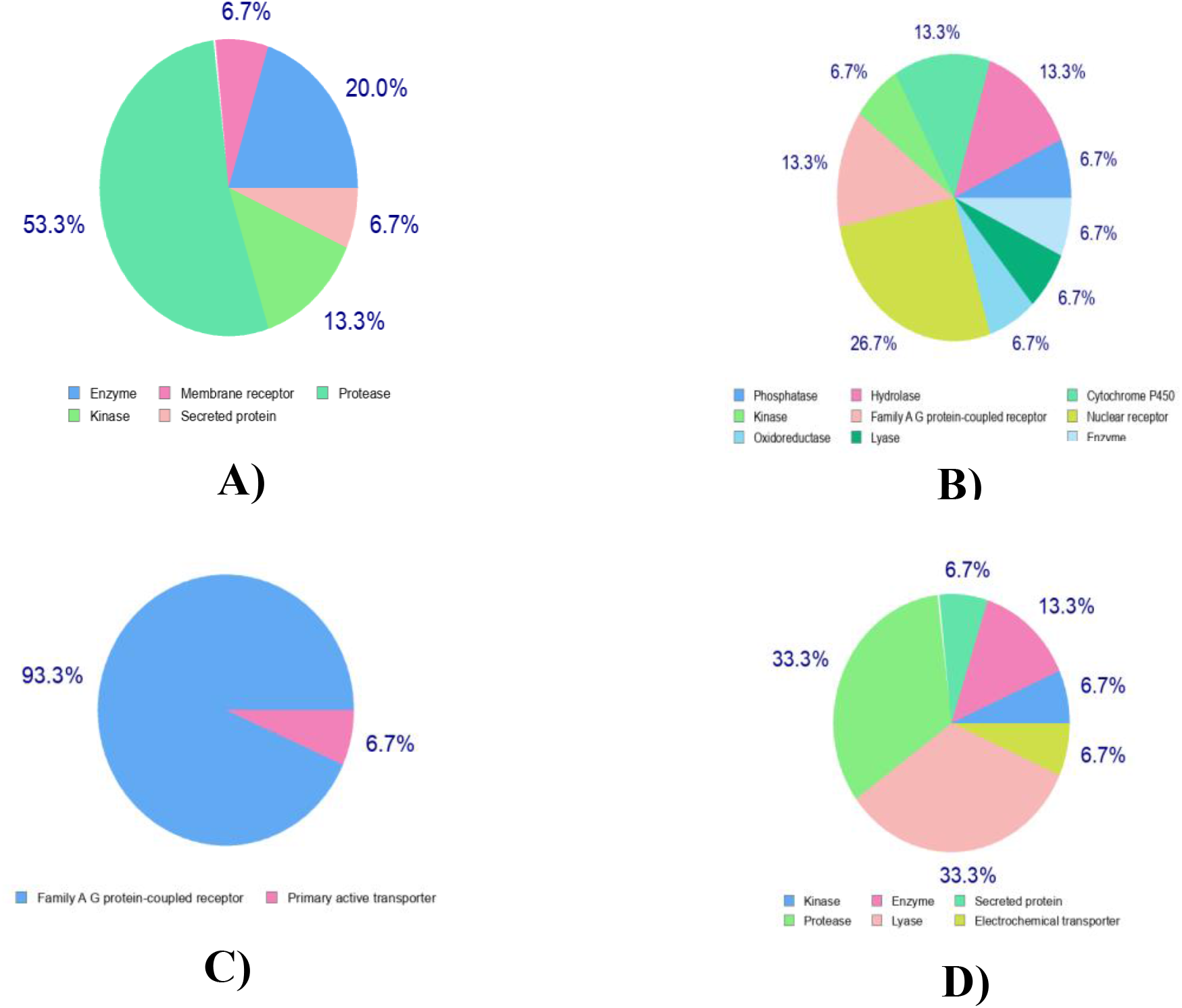
Prediction of drug targets for Chicoric acid (A), Luteone (B), Reserpine (C) and Rosmarinic acid

## Discussion

Before its arrival in the Americas in 2015, the ZIKV has infected millions of people. The virus, primarily due to its connection with serious neurological conditions, has triggered social and sanitary warning (Saiz et al., 2017). Despite of extensive research any vaccine or approved drug has not been released to thwart ZIKA virus infection and reduce the mortality (Panwar and Singh, 2018). By being a lead molecule in the production of drug candidates, plant-derived natural products play an essential role. (Joseph et al., 2016). Plants provide a variety of phytochemicals with antioxidant action, such as flavonoids, terpenoids, lignins, alkaloids and coumarins which provide help to repress viral activity. Therapies based on various plant-derived products for human health are associated with less toxicity and minimal side effects (Ghildiyal et al., 2020). Therefore, attempts have been made in the present analysis to test certain plant-derived metabolites as ZIKA inhibitory agents on the basis of their binding affinities to the pathogen’s several key proteins. In this current investigation, a total of 35 phyto chemicals were tested against four key proteins of the ZIKA virus. The computational biology opens new opportunities and accelerated the drug discovery processes (Hirono 2002). This approach now becomes a core component to discover and develop new lead compounds against many infectious pathogens in the biopharmaceutical industry (Hirono 2002; Ivanov et al., 2006). By this way, the possibilities of binding potential small molecules as ligands / inhibitors can be visualized and save time and cost in the drug development process (Joseph et al., 2016). Plant derived compounds like Luteolin, Quercetin, Baicalein and Kaempferol are potential antiviral agents against a wide range of important viruses including HIV, H5N1 influenza A virus, Dengue virus (DENV), Chikungunya Virus (CHIKV), Japanese encephalitis virus and Coxsackie virus (Habbu et al., 2009). Pinocembrin, a flavanone derived from honey, tea and ruddy wine, is used as a control in present study. It was too appeared that Pinocembrin had broad-spectrum antiviral impacts against ZIKA which is experimentally proved, also shows antiviral activity against DENV2 and CHIKV (Lee et al., 2019).

In this study we evaluate several phyto chemicals against several proteins of ZIKA virus i.e. envelope protein (5JHM), Capsid Protein (5YGH), NS5 RNA-dependent RNA polymerase (5U04), and NS2B-NS3 protease (5LC0) (Dai et al., 2016; Shang et al., 2018; Lei et al., 2016; Godoy et al., 2017).

The NS5 RNA-dependent RNA polymerase (5U04) and NS2B-NS3 protease (5LC0) are the core protein to remain infective and survive for the ZIKA virus (Lei et al., 2016; Godoy et al., 2017). NS2B / NS3 viral protease, which cleaves the viral polyprotein along with host cell proteases into the individual proteins required for viral replication; the NS5 RNA-dependent RNA polymerase of ZIKA also a crucial need for virus replication (Lei et al., 2016; Godoy et al., 2017). The othe two targeted protein, ZIKV-E (5JHM) contains a unique, positively charged patch, which may influence host attachment and the capsid (5YGH) protein is a multifunctional protein, since it binds to viral RNA in the process of nucleocapsid assembly and plays important roles in virus infection processes by interacting with cellular proteins, modulating cellular metabolism, apoptosis and immune response (Dai et al., 2016; Shang et al., 2018). These characteristics make us to work with these four protein and these are an attractive candidate for drug design. Among the plant derived bio active compounds, luteone exhibited the highest binding affinity with Capsid Protein (−52.92 kcal/mol), Chicoric acid with Envelope protein (−58.26 kcal/mol), reserpine with ZIKA virus NS2B-NS3 protease (61.45 kcal/mol) and finally, rosmarinic acid shows best binding affinity with NS5-RdRP (−51.26 kcal/mol) (Figure 2, Table 2). The scores of top candidates were either higher or close some instances than pinocembrin, a positive control used in the present study (Table 2).

Luteone possessed protective activity against fungal infection (Harborne et al., 1976). Reserpine used as an antivral against SARS-CoV virus in europe and USA and also shows Antihypertensive, Antipsychotic properties (Jones 2004 and Baumeisteret al., 2003). Chicoric acid shows antiviral potency against HIV-1 virus (Lin et al., 1999 and Reinke et al., 2004). Rosmarinic acid is effective against Japanese encephalitis virus as antiviral and also shows Antioxidant, Anti-inflammatory phenomenon (Swarup et al., 2007 and Alagawany*et al*., 2017) (Table 1).

Our current findings disclosed the molecular interactions of the top drugs Candidates with key proteins from ZIKA virus (Figure 2, Table 2). The binding modes of our top drugs were simililar with the peptide boronic acid which used as a proteasome inhibitor to determine the crystal structure of the ZIKA protease (Lei et al., 2016) Hydrophobicity is such a large contributor to protein complex stability (Van et al., 2015). The top candidates were well fitted into the active pocket of ZIKA protease protein where several hydrophobic amino acid residues including Met51, Ala1132, Val1155, Gly1133, Pro1131, Val1036 creates a hydrophobic atmosphere, which may help in complexes stabilization. The other protein docked complexes amino acids from 31 to 61 positions were crucial for the binding interactions of ZIKAV Capsid protein (5YGH). Where Capsid protein Arg32, Ala35, Leu33, Lys31 amino acid was involved in most cases to formed the docked complex with phyto-ligands. The ligands shows better binding affinity for 143-145, 301-303 and 361-374 regions for envelop protein (5JHM) and most dominant amino acids for interactions with the ligands were Leu300,Lys301, Val303, Val364, Ala361, Asn362, Thr366, Val143, Gly 145. The protein amino acid 403-423 regions were identified as top surface hotspots for NS5-RdRP (5U04) protein and among binding residues Ala423, Ala408, Lys403 were found most of the cases. In molecular dynamic study, Protein-ligand complex deformability analysis showes that the distortions were lower atomic distortion and higher Eigen values as well which strengthen our statement i.e. complexes shows resistance to deform and remain stable (Figure 4C,5C,6C,7C). The B factor analysis finds a loop region into Envelope-Chicoric acid complex (Figure 4B).The RMSF plot shows minor fluctuations, reflecting the uninterrupted interaction between the protein-ligand complex (Figure 5E,6E,7E). But Envelope-Chicoric acid complex shows a higher bump may be because of a loop region (Figure 4E). In RMSD plot, fluctuations for all complexes were remain within 2 Å (Figure 5D,6D,7D) but Envelope-Chicoric acid fluctuation ranging to 6 Å confirming a bit flexibility (Figure 4D).

ADME results, whether experimentally evaluated or estimated, gives crucial information into how the body actually treats or recognizes a substance. Therefore, although a drug lead can demonstrate phenomenal in vitro effectiveness, weak ADME outcomes almost inevitably terminate its development (Wishart 2007). In predicting potential ADME and toxicity concerns and reducing the number of tests requiring animal experimentation where computational approaches play a key role. To investigate their drug profiles, the top drug applicants were then hired for ADME review. However, none of the metabolites displayed any adverse effects that could decrease their properties of drug likeness. ZIKA often do not attack the brain tissue (Centers for Disease Control and Prevention). Thus, there is no need to permeate the blood brain barrier (BBB) for being an effective molecule against ZIKA. However, no BBB permeates were found among the top drug candidates (Table 3). The ADME analysis shows our top predicted bio active drug candidate has drug likeliness properties (Figure8, Table 3). The toxicity pattern also shows there are no undesirable consequences possessed by these compounds (Table 4). Maximum predicted of the target class for the top drug candidates belonged to the categories of enzymes (e.g. protease, hydrolase, Phosphatase) (Figure 9, Table 5). The similarity analysis displays several similar approved and experimental drug compounds (e.g. Isoformononetin, Deserpidine, Rescinnamine) which may have the potency to cease ZIKA infection (Table 6). The inquiry could be helpful in unraveling the primary drug target hotspot and the medicinal chemistry of the investigational drugs currently being studied against ZIKA.

## Conclusion

The findings indicate that our top predicted bio-active natural drug candidates may be potential inhibitor against ZIKA Virus. Our research findings are promising and for experimental validation we highly recommend further in-vivo trial. This study may also open a door to design a new candidate with better inhibitory activity against notorious ZIKA virus in near future.

## Abbreviations

ZIKV: ZIKA virus
LD_50_: Lethal dose 50
ADME: Absorption, distribution, metabolism, and excretion
BBB: Blood brain barrier.

## Acknowledgements

Authors would like to acknowledge the Department of Plant and Environmental Biotechnology of Sylhet Agricultural University, Sylhet-3100, Bangladesh for the technical support of this research.

